# Short communication: Efficacy of two commercial disinfectants on *Paenibacillus larvae* spores

**DOI:** 10.1101/2021.03.16.435711

**Authors:** Joseph Kiriamburi, Jamleck Muturi, Julius Mugweru, Eva Forsgren, Anna Nilsson

**Affiliations:** Department of Biological Sciences, University of Embu, P.O Box 6 -60100, Embu - Kenya; Department of Ecology, Swedish University of Agricultural Sciences, PO Box 7044, Uppsala 750 07, Sweden

**Keywords:** *Apis mellifera*, American foulbrood, *Paenibacillus larvae*, disinfectants

## Abstract

*Paenibacillus larvae* is a spore-forming bacterium causing American foulbrood (AFB) in honey bee larvae. The remains of a diseased larva contains billions of extremely resilient *P. larvae* spores viable for decades. Burning clinically symptomatic colonies is widely considered the only workable strategy to prevent further spread of the disease, and the management practices used for decontamination requires high concentrations of chemicals or special equipment.

The aim of this study was to test and compare the biocidal effect of two commercially available disinfectants, “ Disinfection for beekeeping” and Virkon S on *P. larvae*. The two products were applied to *P. larvae* spores in suspension as well as inoculated on two common beehive materials, wood and styrofoam.

“ Disinfection for beekeeping” had a 100 % biocidal effect on *P. larvae* spores in suspension compared to 87.0-88.6 % for Virkon S which, however, had a significantly better effect on *P. larvae* on styrofoam. The two disinfectants had similar effect on infected wood material.

## INTRODUCTION

*Paenibacillus larvae* is a spore-forming, Gram-positive bacterium causing the severe disease American foulbrood (AFB) in honey bee larvae. Honey bee larvae become infected from ingesting food contaminated with *P. larvae* spores that germinate in the midgut and eventually kills the larvae. The remains of the larvae contains billions of spores and serves as sources for new infections. The *P. larvae* spores are resilient and can remain viable in the environment for decades (Dobbelaere et al., 2001; Forsgren et al., 2008; Haseman, 1961). A common way to control AFB is by burning the contaminated hives and bees, although the latter can sometimes be saved as an artificial swarm, housed on new or disinfected material (Genersch, 2010). Hive material can be decontaminated using chemical disinfectants or heat. Chemical disinfectants have been shown to have a high efficacy on spores in suspension, but less effective on wood-based equipment (Okayama et al., 1997; Dobbelaere et al., 2001). There are several methods using heat for decontamination of hive material, for example dipping in hot paraffin, scorching, dry heat and autoclaving (Dobbelaere et al., 2001). These methods are effective (Del Hoyo et al., 1998; Dobbelaere et al., 2001), but requires access to advanced equipment.

Our aim was to test and compare the biocidal effect of 2 disinfectants, “ Disinfection for beekeeping” (DFB) (Swienty, Denmark) and Virkon S (Lanxess, Germany) on *P. larvae* spores. DFB is developed for disinfection of hive material, gloves and tools and, according to the manufacturer (www.swienty.com, viewed October 9 2019), have a 99.99% biocidal effect on all viruses, bacteria, spores and fungi. Virkon S is a common disinfectant that have been on the market for over 30 years, originally developed for farm and livestock production (Hernández et al., 2000) (www.virkons.se, viewed October 8 2019).

## MATERIAL AND METHODS

A spore suspension was prepared from *P. larvae* cultures on agarplates (14 days to obtain sporulation) in sterile 0.9% saline solution. The spore suspension was stored at 4°C, heat shocked at 85°C for 10 minutes and diluted to the desired concentrations before the start of each experiment.

The experiments were performed as described in Fig. 1 and repeated at least 3 times. *P. larvae* were cultured according to standard cultivation methods (Nordström & Fries, 1995).

**Fig. 1.**
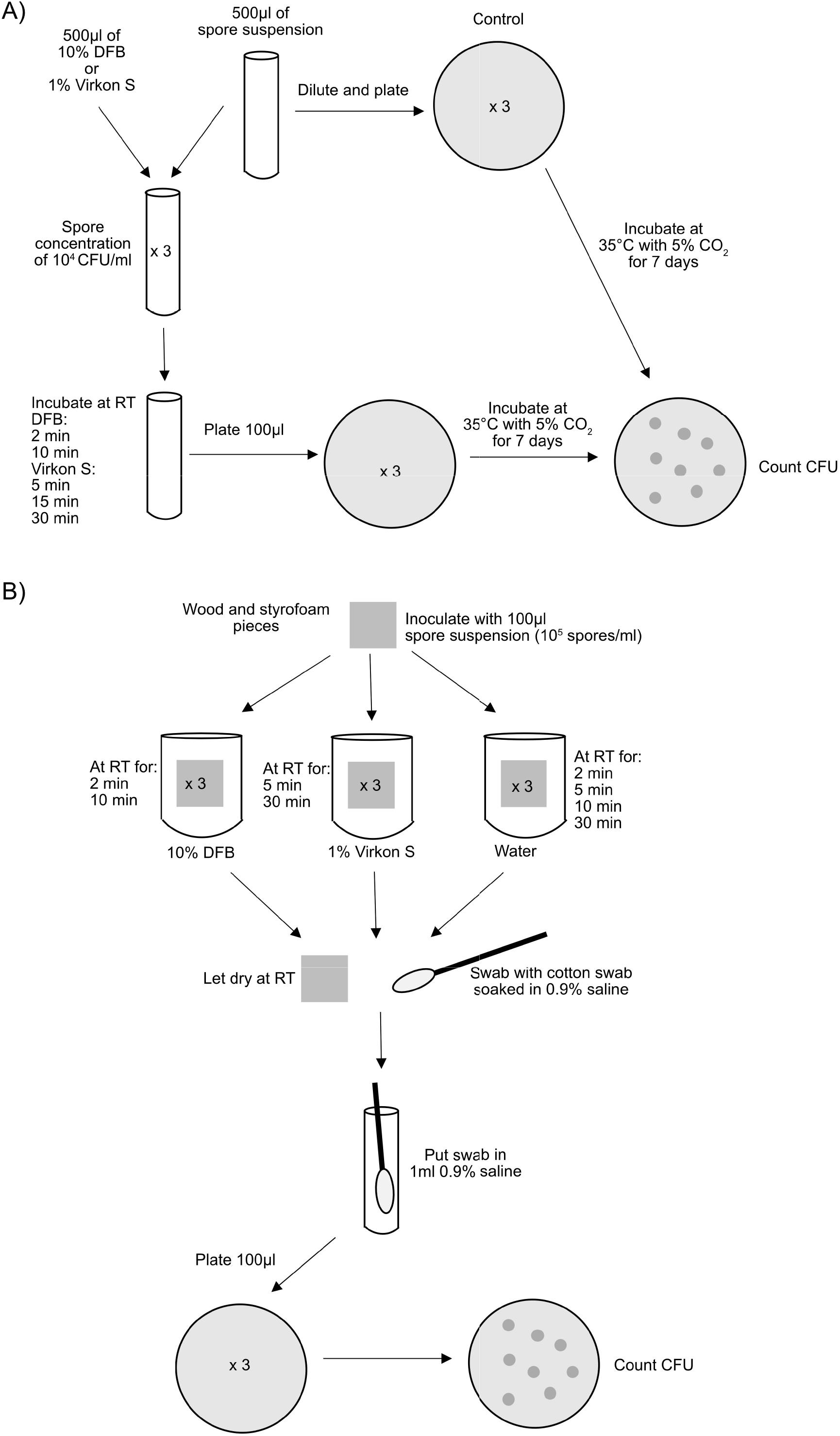
Schematic description of the experimental setup. RT, room temperature.

The biocidal effect of the disinfectants was calculated by comparing the number of CFUs from the treated samples and the untreated spore suspension or the mock treated wood and styrofoam pieces.

Student’s t test (unpaired, 2-tailed) was used to identify statistically significant differences, with a P-value of ≤0.05 considered significant.

## RESULTS

DFB had the highest biocidal effect (100% already after 2 minutes) on spores in suspension and was significantly more efficient than the 5 and 15 minute Virkon S treatments (all P=0.01, fig. 1A).

On wood, no significant differences could be seen between DFB and Virkon S, or the between the different treatment times (fig. 1B).

On Styrofoam, a significantly higher biocidal effect was observed after 30 minutes treatment with Virkon S compared to 2 and 10 minutes treatment with DFB (both P=0.02, fig. 1C). The 30 minute treatment with Virkon S had also a significantly higher biocidal effect than the 5 minute treatment (P=0.01, fig. 1C).

## DISCUSSION

This study compares the biocidal effect of 2 disinfectants on *P. larvae* spores. Both disinfectants had an effect on the bacterial spores in suspension and on wood and styrofoam. DFB had the best effect on the bacterial spores in suspension where all *P. larvae* spores were killed. These results are in line with the information from the manufacturer saying that DFB kills all viruses, bacteria, fungi and spores within 45 s. However, the effect of DFB on spores on wood and styrofoam was lower than in suspension (fig. 1). Virkon S was slightly less effective than DFB on spores in suspension, but the differences were not significant. Thirty minutes treatment (recommended by the manufacturer) of Virkon S on contaminated styrofoam was significantly more effective than the treatment with DFB (fig. 1C). Virkon S has in a previous study been shown to kill 80% of *P. larvae* spores (Hansen & Brødsgaard, 1999). In this study however, the biocidal effect ranged from 88.6 to 96.8% after 30 minutes treatment (fig. 1). The effect of both disinfectants on wood varied more than the effect on styrofoam and in suspension, most likely due to difficulties recovering *P. larvae* from wood. This is probably because wood is more porous and absorbs the liquid with the spores. *P. larvae* spores can “ hide” in wood, making it more difficult for the disinfectant to access the bacterium. The wood and styrofoam pieces used in this study were clean, *i*.*e*. they were not covered in wax or propolis. Any disinfectants will probably be less effective on used, non-cleaned hive material where large amounts of bacterial spores may be inaccessible to the disinfectants. It is therefore important that infected materials are thoroughly cleaned before being treated with disinfectants.

## ACKNOWLEDGEMENTS

We thank Karin Ullman for technical assistance and Preben Kristiansen for valuable comments, and we are thankful for the support from the Linnaeus-Palme international exchange program.

---

**Fig. 2.**
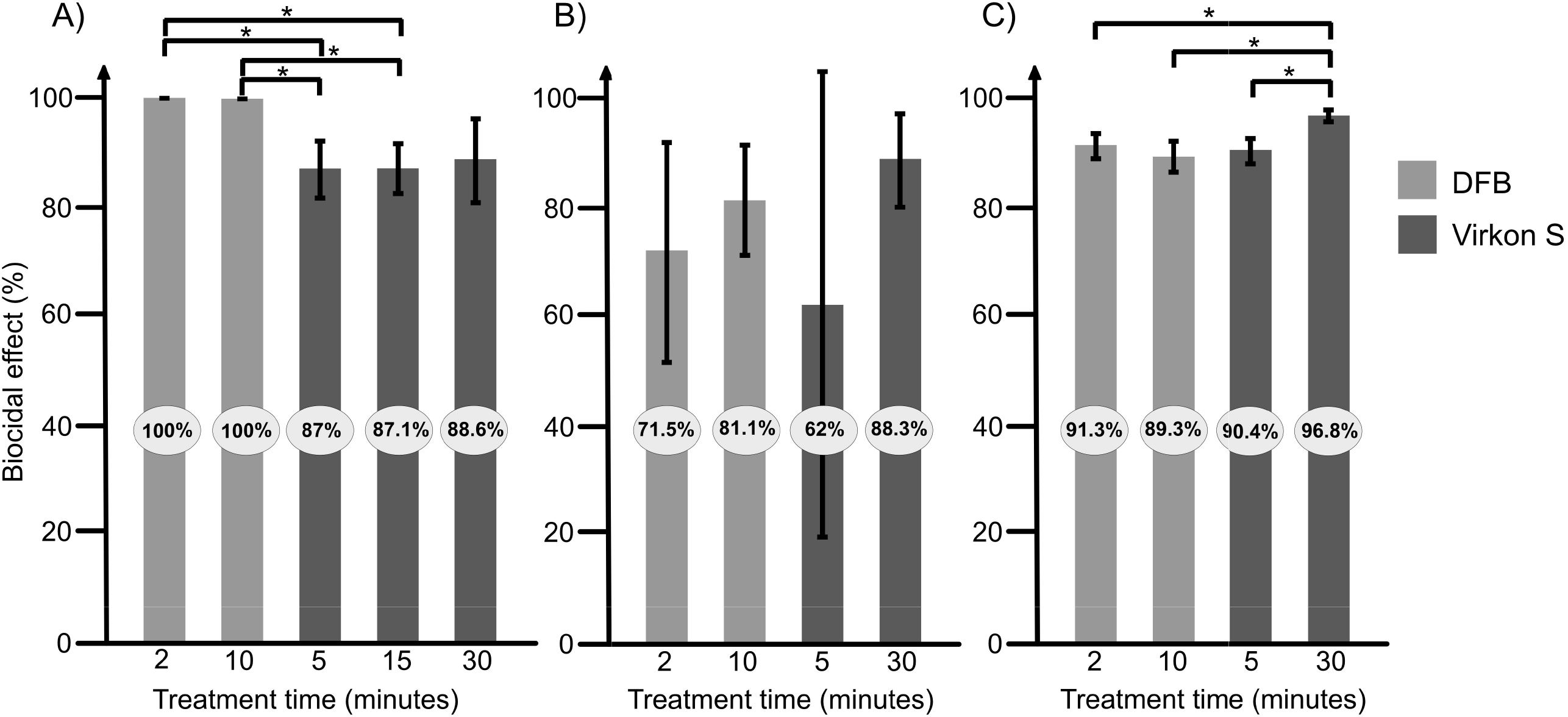
Efficacy of the 2 disinfectants “ Disinfection for beekeeping” (DFB) and Virkon S on *P. larvae* spores in suspension (A), on wood (B) and on styrofoam (C). Light grey bars shows results for DFB and darker grey Virkon S. The result is presented as an average from at least 3 repeats, with error bars indicating standard deviation. Significant differences are indicated. *, P<0.05.

## References

Del Hoyo, M., Basualdo, M., Torres, J., & Bedascarrasbure, E. (1998). Use of DHT-Equpment for Disinfection of AFB-Contaminated Beehive Materials in Argentina. American Bee Journal, 138(10), 738–740.

Dobbelaere, W. et al. (2001). Disinfection of wooden structures contaminated with Paenibacillus larvae subsp. Larvae spores. Journal of Applied Microbiology, 91(2), 212–216.

Forsgren, E., Stevanovic, J., & Fries, I. (2008). Variability in germination and in temperature and storage resistance among Paenibacillus larvae genotypes. Veterinary Microbiology, 129(3), 342–349.

Genersch, E. (2010). American Foulbrood in honeybees and its causative agent, Paenibacillus larvae. Journal of Invertebrate Pathology, 103, S10–S19.

Hansen, H., & Brødsgaard, C. J. (1999). American foulbrood: A review of its biology, diagnosis and control. Bee World, 80(1), 5–23.

Haseman, L. (1961). How Long Can Spores of American Foulbrood Live? American Bee Journal, 101, 298–299.

Hernández, A. et al. (2000). Assessment of in-vitro efficacy of 1% Virkon® against bacteria, fungi, viruses and spores by means of AFNOR guidelines. Journal of Hospital Infection, 46(3), 203–209.

Nordström, S., & Fries, I. (1995). A comparison of media and cultural conditions for identification of Bacillus larvae in honey. Journal of Apicultural Research, 34(2), 97– 103.

Okayama, A. et al. (1997). Sporicidal Activities of Disinfectants on Paenibacillus larvae. Journal of Veterinary Medical Science, 59(10), 953–954.

